# GPseudoRank: a permutation sampler for single cell orderings

**DOI:** 10.1101/211417

**Authors:** Magdalena E Strauß, John E Reid, Lorenz Wernisch

## Abstract

**Motivation:** A number of pseudotime methods have provided point estimates of the ordering of cells for scRNA-seq data. A still limited number of methods also model the uncertainty of the pseudotime estimate. However, there is still a need for a method to sample from complicated and multi-modal distributions of orders, and to estimate changes in the amount of the uncertainty of the order during the course of a biological development, as this can support the selection of suitable cells for the clustering of genes or for network inference.

**Results:** In an application to a microarray data set our proposed method, GPseudoRank, identifies two modes of the distribution, each of them corresponding to point estimates of orders obtained by a different established method. In an application to scRNA-seq data we demonstrate the potential of GPseudoRank to identify phases of lower and higher pseudotime uncertainty during a biological process. GPseudoRank also correctly identifies cells precocious in their antiviral response.

**Availability and implementation:** Our method is available on github: https://github.com/magStra/GPseudoRank.

**Supplementary information:** Supplementary materials are available.

## 1 Introduction

Providing mRNA expression levels of genes for individual cells, scRNA-seq has shown heterogeneity of gene expression across cells during various biological developments. While part of this results from technical noise, part is generally attributable to genuine cell heterogeneity. See, for instance, [5, 31]. Due to the destruction of the cells as a result of the measurement process, scRNA-seq only provides a single measurement per cell [28], never time series data following the development of the same single cell. However, individual cells progress through changes at different time scales [30]. Thus it is possible to obtain a form of time series data even from cross-sectional data by statistical means, an approach referred to as pseudotime ordering.

Most approaches to pseudotemporal ordering are based on representing cells as *n*_*g*_-dimensional vectors, where *n*_*g*_ is a selected number of genes in a cell. Algorithms exploit the neighborhood structure of these vectors to find a pseudotemporal ordering, a linear ordering of all or most cells so that cells which are close in ℝ^*n_g_*^ are also close in the linear ordering.

Wanderlust [4] and SLICER [32, 33] are two examples of methods based on *k* nearest neighbours graphs. SLICER additionally first applies LLE (local linear embedding) [25] for dimensionality reduction. A number of methods are based on diffusion maps [3, 11, 12, 26]. TSCAN [16, 15] is based on the construction of a minimum spanning tree (MST) between centroids of clusters, with an intermediate clustering step. Another well-known method using MST and clustering is Monocle 2 [22], which applies graph structure learning [17].

The approaches mentioned above and a number of others provide singular pseudotime orderings without modelling uncertainty. Campbell and Yau [7] examined the stability of Monocle’s pseudotime estimation when applied to random subsets of cells. They showed that the estimates can vary significantly. Thus quantification of uncertainty in pseudotime is crucial to avoid overconfidence. There are two existing methods for pseudotime estimation using MCMC to sample from a posterior distribution [7, 24], and a few others using variational methods [1, 24, 34]. They use Gaussian processes (GPs, see Section 2.1) to model the data. However, these methods sample from, or approximate, in the case of variational inference, posterior distributions of continuous pseudotime vectors in ℝ^*n*^, rather than sampling the ordering as a permutation.

We propose GPseudoRank, an algorithm sampling from a posterior distribution of pseudo-orders instead of pseudotimes, avoiding the exploration of pseudotime assignments that all map to the same ordering. MCMC samplers (such as NUTS [14]) suitable for use in continuous pseudotime spaces make local moves that can have problems exploring bi-modal posteriors. GPseudoRank, by contrast, exploits a range of local and long-distance MCMC moves tailored to efficiently traverse the space of permutations. It also provides continuous pseudotime estimates by deriving a pseudotime vector from a fixed ordering through a deterministic transformation. This is based on the observation that most continuous pseudotime vectors with high likelihood are concentrated around pseudotime vectors derived from orderings through this transformation.

## 2 Methods

### 2.1 Single-cell trajectories as stochastic processes

We assume we have preprocessed logarithmised gene expression data in the form *y*_*g*_ (*c*) of gene *g*, *g* = 1,…, *n*_*g*_, in cell *c*, *c* = 1,…, *T* (see section 2.6 for preprocessing steps). We start with a vector of time points ***τ*** = (*τ*_1_, …, *τ_T_*) and define an ordering of cells as a permutation **o** = (*o*_1_,…, *o*_*T*_), *o_i_* ∈ {1,…, *T*}, *o_i_* ≠ *o_j_* for *i* ≠ *j*, where *o_i_* is the index of the cell assigned to time *τ_i_*. We model the gene expression trajectories *y*_*g*_ = (*y*_*g*_(*o*_1_),…,*y*_*g*_(*o*_*T*_)) for each gene *g* by Gaussian processes (GPs) [23], conditional on an ordering **o** of the cells. A GP is a distribution over functions of time in terms of a mean function *μ* and a covariance function Σ. For an input vector ***τ*** = (*τ*_1_,…,*τ_T_*) of time points, *μ*(***τ***) returns a vector of *T* mean values of function evaluations at these time points and Σ(***τ***) a *T* × *T* matrix of covariances of function evaluations at the time points. The distribution of functions *f* ~ *GP*(***μ***, **Σ**) is described by stating that, for any vector of time points ***τ*** = (*τ*_1_,…,*τ_T_*), evaluations *f*(*τ_i_*) follow a multivariate normal 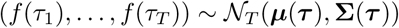. In this study we use a squared exponential covariance function for Σ.

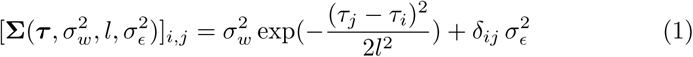

where 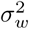 is a scale parameter, *l* a length scale and 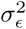 a term representing measurement noise.

Given an ordering **o**, the expression data for gene *g* can be ordered accordingly: *y*_*g*_(**o**) = (*y*_*g*_(*o*_1_),…,*y*_*g*_(*o*_*T*_)) and we model this trajectory as

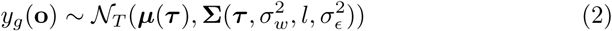

for each gene *g* = 1,…,*n*_*g*_, where ***τ*** = (*τ*_1_,…,*τ_T_*) are time points. In practice, we assume a zero-mean GP, that is, ***μ*** = 0. To adjust the data for this assumption we subtract the overall mean across all genes and cells from each entry in the matrix of gene expression levels (see Sections 2.6.2 and 2.6.3).

### 2.2 Geodesic mapping

Pseudotime should not be confused with physical time in which cell development unfolds. In order to identify the latent time points ***τ*** = (*τ*_1_,…,*τ_T_*), which we assume to be unknown, together with the smoothness parameters of the GP, we have to make additional assumptions. The overall scale can be fixed by assuming *τ_i_* ∈ [0,1] and each cell could be assigned some *rank time*, equidistant time points ((*i* − 0.5)/*T* | *i* = 1,…,*T*). Rank time is similar to the concept of master time developed in [34]. However, rank time depends on the number of cells sampled per capture time, which could be rather arbitrary, and does not allow for any local change in scale. We therefore suggest a different route to identify latent time points. We assume the covariance structure, essentially the smoothness of the process, is independent of time, that the GP is *stationary*. Pseudotime can then be considered a latent variable measuring biological development rather than physical time [1, 7, 24, 34]. For periods of slower development, for example, pseudotime intervals will be shorter than physical time intervals and longer for faster development. In order to account for such change in scale over time we compute time points for any given ordering **o** as follows (recall *o_j_* is the index of the cell in position *j*).

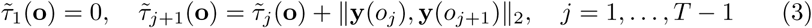

where **y**(*o_j_*) = (*y*_1_(*o_j_*),…,*y_n_g__*(*o_j_*))^*T*^ and ∥·∥_2_ is the Euclidean norm in ℝ^*n_g_*^. Then we set 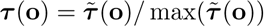 to obtain pseudotimes ***τ***(**o**) in the interval [0,1]. For cells next to each other in the order **o**, this mapping puts them closer in pseudotime if they are similar in their expression profiles and further apart if they are less so. That is, the *j*-th time point *τ_j_* is the geodesic distance of cell *o_j_* from the first cell *o*_1_, where we approximate the geodesic distance as the sum of the Euclidean distances between the cells ranked next to each other, similar to the dimensionality reduction method Isomap [29]. Geodesic distances have previously been used for pseudotime estimation, see for instance [22, 32]. The importance of allowing pseudotime to deviate from rank time for sampling from the correct posterior distribution is illustrated in more detail in Section 3.2 below and Section 2 of the supplementary materials.

### 2.3 Gaussian process priors

The correct ordering **o** of cells is distinguished by comparatively low measurement noise 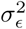 in (1), since most of the variation is captured by the trajectory whose variability is determined by the scale parameter 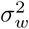. Therefore informative priors for the noise parameters are necessary to ensure the model concentrates probability mass around the correct order and to avoid that a sampling or estimation algorithm gets trapped in local modes. Furthermore, since total variability is a sum of measurement noise and signal variability, we sample only 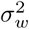 and set 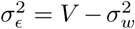, where *V* is the sample variance taken across the entire *n_g_* × *T* matrix of gene expression levels of *T* cells for *n_g_* genes. The priors are as follows:

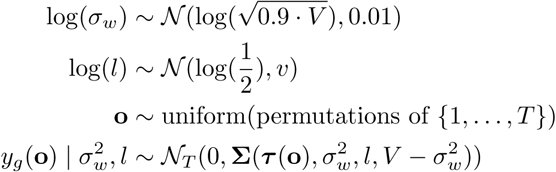

We set *v* = 0.1 for the microarray data set considered in [35] (see Section 2.6.2) and *v* = 0.01 for the scRNA-seq data set [27] (see Section 2.6.3).

### 2.4 MCMC sampling

Markov Chain Monte Carlo (MCMC) methods [10] have been widely used to sample from continuous posterior densities in Bayesian statistics. They construct Markov Chains with the posterior distribution as their equilibrium. After convergence, each sample from the MCMC is taken as a sample from the posterior distribution. Our proposed method uses the Metropolis-Hastings algorithm [13, 18] for the sampling. For each given state of the Markov Chain, a new state is proposed using a proposal distribution, and accepted if an acceptance ratio is less than a uniform random number. While the construction of proposal distributions is often straightforward in the continuous case, we developed novel proposal moves to sample from discrete distributions of orders (see Section 2.5). For the sampling of the GP parameters we use Gaussian proposal distributions, adapting their standard deviation during burn-in aiming at acceptance rates between 0.45 and 0.5.

### 2.5 Sampling orderings

In the following we propose a Metropolis-Hastings algorithm for the sampling of the orderings. Preliminary experience with a variety of combinatorial moves to sample permutations led to the following set of five core moves, each with probability *p_j_*, *j* = 1,…,5:

1. Move 1, **iterated swapping of neighbouring cells**: draw the number *r*_1_ of swaps to be applied uniformly from 1,…,*n*_0_ and draw *r*_1_ swap positions *P*_1_,…,*P*_*r*_1__ from 1,…,*T*−1 with replacement. Then iterate for *j* = 1,…,*r*_1_: swap cell at position *P_j_* with its neighbor at position *P_j_* + 1.
2. Move 2, **swapping of cells with short *L*^1^-distances**: select two positions *i* and *j* according to probability *p_ij_* ∝ exp(−*d*(*c_i_*, *c_j_*)^2^/*γ*_1_), where *d* refers to the *L*_1_ distances of cells *c_i_* and *c_j_* (as *n_g_*-dimensional vectors) in these positions. Move *c_i_* to position *j* and *c_j_* to position *i*.
3. Move 3, **reversing segments between cells with short *L*^1^-distances**: obtain two positions *i* and *j* as in move 2 and reverse the ordering of all cells in between, including cells at *i* and *j*.
4. Move 4, **short random permutations**: draw a number *r*_2_ of short permutations uniformly from 1,…,*n*_3_. For each *j* = 1,…,*r*_2_, draw a number *r*_3,*j*_ uniformly from 3,…,max(*n*_3*a*_, 3)) and a cell position *k_j_* uniformly from 1,…,*T*−*r*_3,*j*_. Randomly permute the cells at positions *k_j_*,…,*k_j_*+*r*_3,*j*_.
5. Move 5, **reversing the entire ordering**.

The rationale for moves 2 and 3 is that two cells which are positioned apart in the ordering should only be exchanged (move 2) or the segment between them reversed (move 3) if these cells have similar expression profiles and the smoothness of the trajectory remains intact after the move. For move 1 we use a default setting of *n*_0_ = [*T*/4] for the simulation studies. For move 4 we set *n*_3_ = [*T*/20], and *n*_3*a*_ = [*T*/12]. The distributions for choosing moves 2 and 3 may be tempered, that is taken to the power of a factor 0 < *α* < 1, to lower acceptance rates if required.

For the simulation studies we apply all possible combinations of moves 1 to 4 with equal probabilities and move 5 with a probability of 0.002. For the microarray data we apply only move 3, as (as will be shown below) it is the best sampling strategy for multi-modal distributions. For the scRNA-seq data set, we use moves 1 to 4 with probability 0.2495, and move 5 with probability 0.002. For the microarray data set we use *γ* = 1000 in move 3 and an additional tempering factor *a* = 0.1. For the scRNA-seq data set we set *γ* = 4000, without any tempering factor for moves 2 or 3.

As our posterior distribution is a symmetric function of the order, each order and its reverse will be sampled with equal probability from the posterior distribution. We remove this symmetry in further analysis by reversing orders which are negatively correlated with the capture times.

### 2.6 Data sets

#### 2.6.1 Simulated data

The efficacy of the individual moves and of combinations of different moves for different types of data is first assessed on simulated data. We simulate *n*_*g*_ = 50 genes for *T* = 90 cells. For each simulation study we generate 16 data sets. On each of these data sets we run MCMC chains using all the possible combinations of the four proposed moves (with equal probability for combinations of more than one move). Since in the simulations we are mostly interested in the assessment of ordering moves and not any parameter estimation, we fix them to their true values and fix time points to rank time.

**Simulation 1: three capture times, low noise**. Each of the 16 data sets is generated as follows. First 90 temporal input points are drawn uniformly from [0,1]. Then for each of the 50 genes in each of the simulated data sets, a parameter set for a GP underlying the trajectory of the simulated gene is drawn from

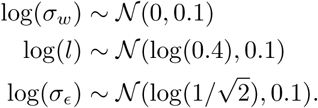

The data are assumed to be obtained at three capture times with 30 cells each.

**Simulation 2: two capture times, low noise**. The setup is similar to simulation 1, but with two capture times, where 30 cells are assigned to the first capture time, and the remaining 60 to the second.

**Simulation 3: three capture times, high noise**. The setup is similar to simulation 1, but 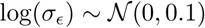.

#### 2.6.2 Microarray data

Windram et al. [35] studied the response of Arabidopsis thaliana to infection by the fungal pathogen Botrytis cinerea, generating microarray time series data over 48 hours, with measurements at intervals of 2 hours. As in Reid and Wernisch [24], we assume 4 capture times with 6 cells each. We compare the result to estimates produced by two established pseudotime methods, TSCAN [16, 15] and SLICER [32, 33], using the standard settings for the latter algorithms. For SLICER setting the number of edges of the nearest neighbours graph in the low dimensional space to 4 and 5 resulted in orders closest to the true one. For all analyses, we use the 150 genes mentioned in the paper by Windram et al. [35].

#### 2.6.3 Single cell RNA-seq data

Shalek et al. [27] examined the response of primary mouse bone-marrow-derived dendritic cells in three different conditions using single-cell RNA-seq. We apply GPseudoRank to the lipopolysaccharide stimulated (LPS) condition. Shalek et al. [27] identified four modules of genes. As in Reid and Wernisch [24], we use a total of 74 genes from the four modules with the highest temporal variance relative to their noise levels [24]. The number of cells is 307, with 49 unstimulated cells, 75 captured after 1h, 65 after 2h, 60 after 4h, and 58 after 6h. We use an adjustment for cell size developed by Anders and Huber [2], also used in Reid and Wernisch [24].

### 2.7 Convergence assessment

For thorough convergence assessment, we run 12 different chains for each of the real data sets, and 5 for each of the simulation set-ups. For the simulated and the microarray data sets we run 100,000 iterations per MCMC chain and apply a thinning factor of 10. For the scRNA-seq data we use the same thinning factor, but 500,000 iterations. In order to assess convergence and not to bias the sampler towards specific orderings, all chains are seeded with random starting orders and with random GP parameters sampled from the prior distribution. However, we do restrict starting orders to permutations of cells within, but not across capture times.

To check convergence, we use the Gelman-Rubin 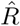-statistic [8], corrected for sampling variability [6], implemented in the R-package coda [21]. The 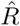-statistic estimates the factor by which the pooled variance across all the chains is larger than the within-sample variance. For convergent chains, 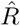 approaches 1 as the number of samples tends to infinity. According to [6], convergence may be assumed to have been reached if 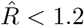. We apply the stricter recommendation of 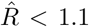 [9]. We compute the 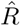 statistics for the following two quantities: first, the log-likelihood, and second the *L*^1^-distances of the sampled cell positions from a fixed reference set of cell positions, for which we use the true order, if known, and 1,…,*T*, where *T* is the number of cells, in case of scRNA-seq data. We compute the 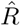 statistics a number of times during sampling, each time discarding the first 50% [9]. We compare the speed of convergence for different combinations of proposal moves in the simulation studies. See Section 1 in the supplementary materials for details.

## 3 Results

### 3.1 Simulation studies

This section summarises the insights gained from the simulation studies. For details on thassessment criteria and results, see Section 1 in the supplementary materials.

**Simulation 1**. Any combination of moves leads to good convergence, and although there are differences in the speed and level of convergence, any combination of moves is recommended.

**Simulation 2**. There are only two capture times, hence there is more variety in the starting orders for each chain. The performance of the combinations of moves is different from simulation 1. Move 3 performs better than any other single move.

Move 3 generally traverses the space of permutations faster by reversing whole segments of an ordering and it is the only move for which all 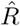-statistics go below 1.1 within the first 10,000 thinned samples. The combination of moves ranked first according to the criteria described in Section 1 of the supplementary materials is the combination 1,2,3,4 of all the moves.

**Simulation 3**. All moves and combinations thereof perform well in this situation, though move 3, while still achieving reasonable levels of convergence, is now the comparatively less well performing single move. The combination of all four moves performs well.

### 3.2 Validation on microarray data

The experimental data set has been acquired at equidistant time points every 2 hours. However, to adjust for differences in the speed of biological development during the process, we apply GPseudoRank with irregular pseudotimes (as explained above in Section 2.2). In fact, adjusting for the speed of biological development is needed, and an approximation with simple equidistant input points for the GP changes the posterior distribution significantly, as shown in Section 2 of the supplementary materials. Because of the bi-modality of the *L*^1^-distances from the true cell positions (Figure 2), the Gelman-Rubin statistic for this distance is less useful and we show the statistic for the log-likelihood instead (Figure 1).

**Figure 1:**
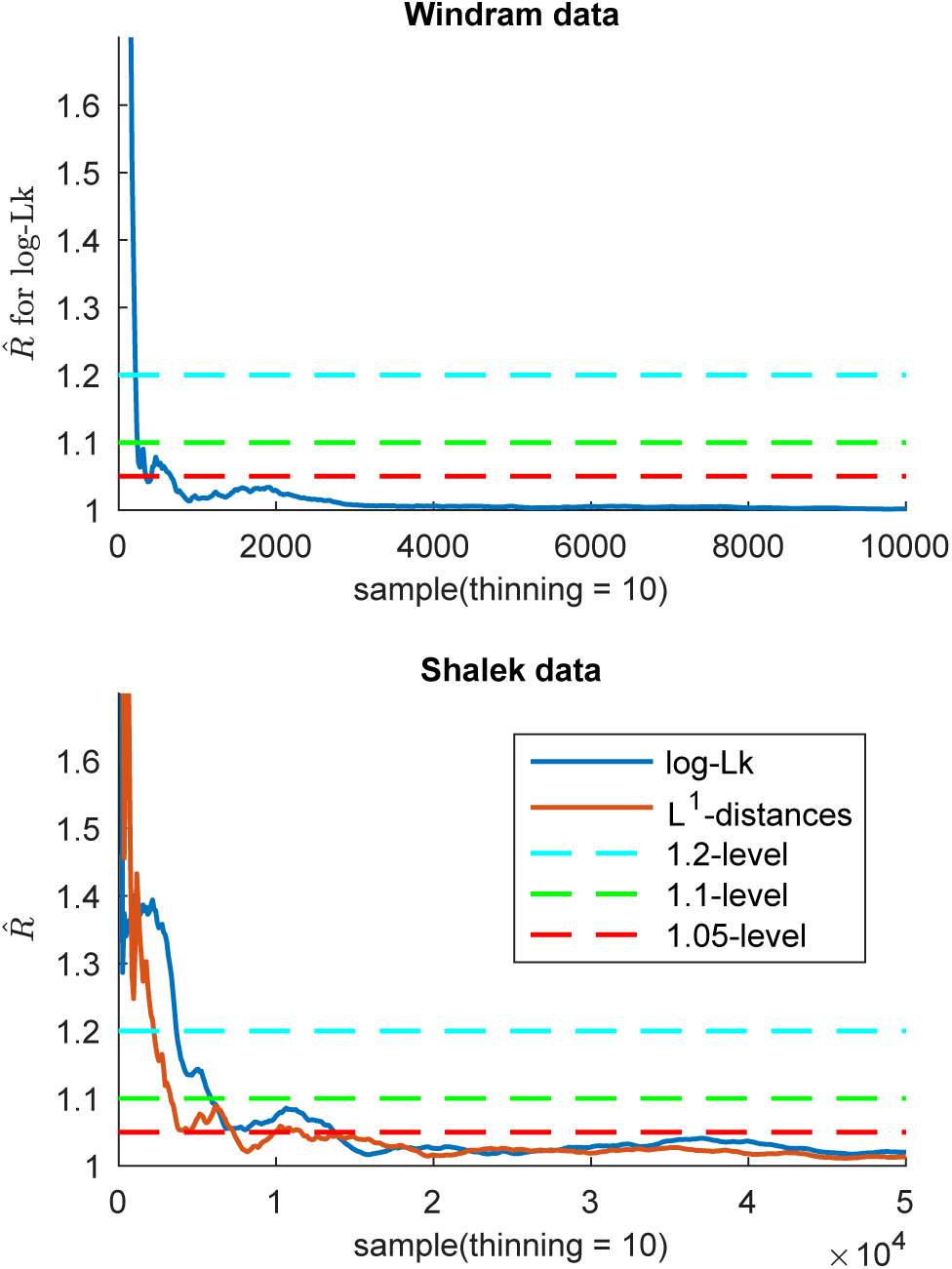
Convergence analysis for GPseudoRank. Gelman-Rubin statistics for the log-likelihood and for the *L*^1^-distances of the sampled permutations of cell positions from the reference permutation (Shalek data).

**Figure 2:**
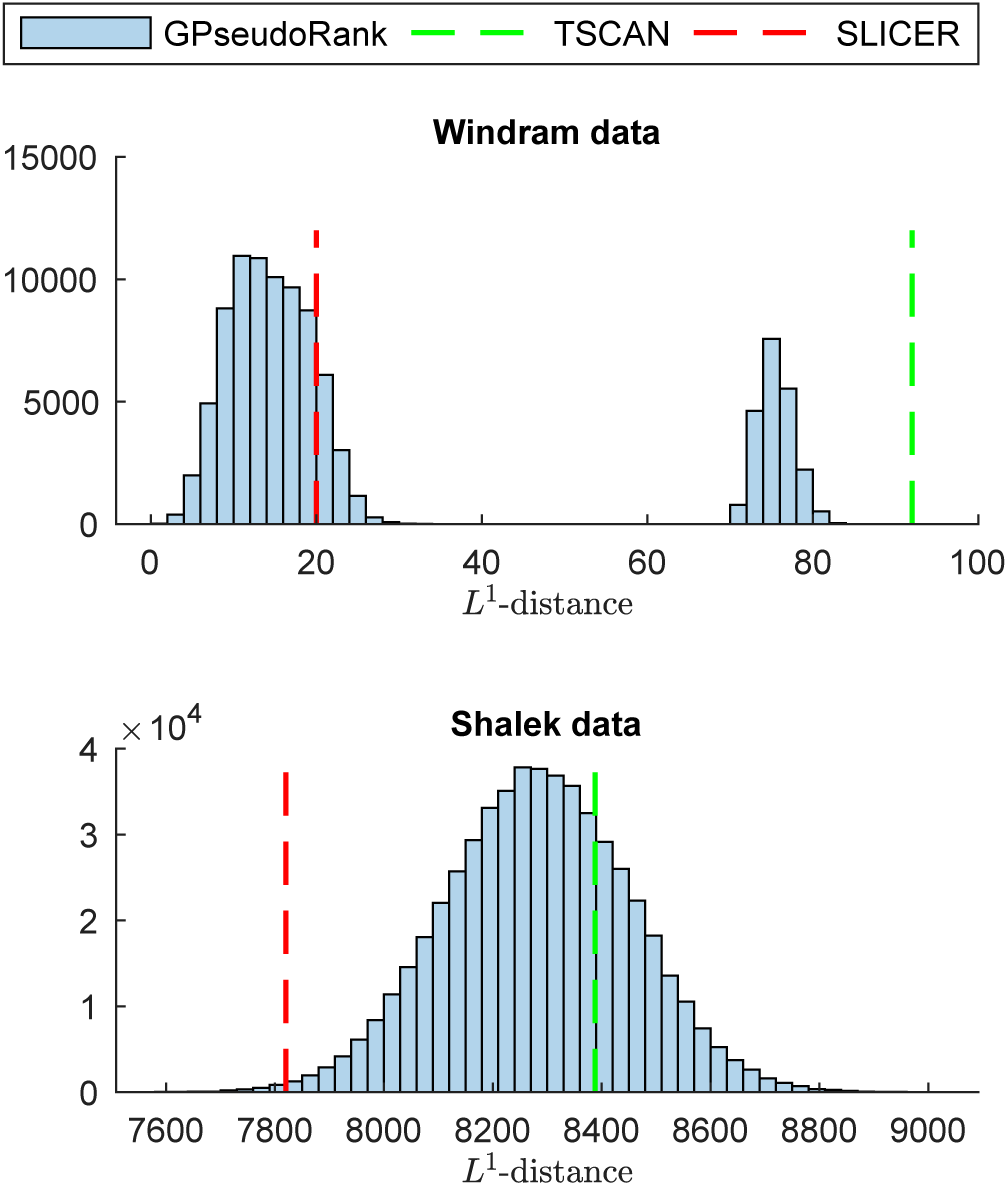
Histogram of L^1^-distances from the reference permutation of cell positions. Distribution sampled with GPseudoRank, point estimates with TSCAN and SLICER.

Figure 1 suggests a very fast convergence. However, if multi-modality is suspected, which might not show in the log likelihood trace, sampling beyond convergence for the log likelihood is recommended. Further plots illustrating convergence can be found in Section 1 of the supplementary materials.

As illustrated by Figure 2, the distribution of the *L*^1^-distances of the sampled permutations of cell positions from the correct order is bi-modal. The estimates provided by SLICER [32, 33] and TSCAN [16, 15] fall in different modes illustrating the importance of sampling from the distribution of the orderings rather than just obtaining a single estimate. The DeLorean MCMC sampler, sampling continuous pseudotimes, is also unable to capture the multi-modality of the posterior [24, Figure 1]. Figure 3 illustrates that the posterior distribution sampled by GPseudoRank covers the two point estimates obtained from TSCAN and from SLICER. For GPseudoRank, it contains the samples from one randomly selected MCMC chain. For the corresponding plots of each of the 12 chains for convergence analysis, see Section 2 in the supplementary materials.

**Figure 3:**
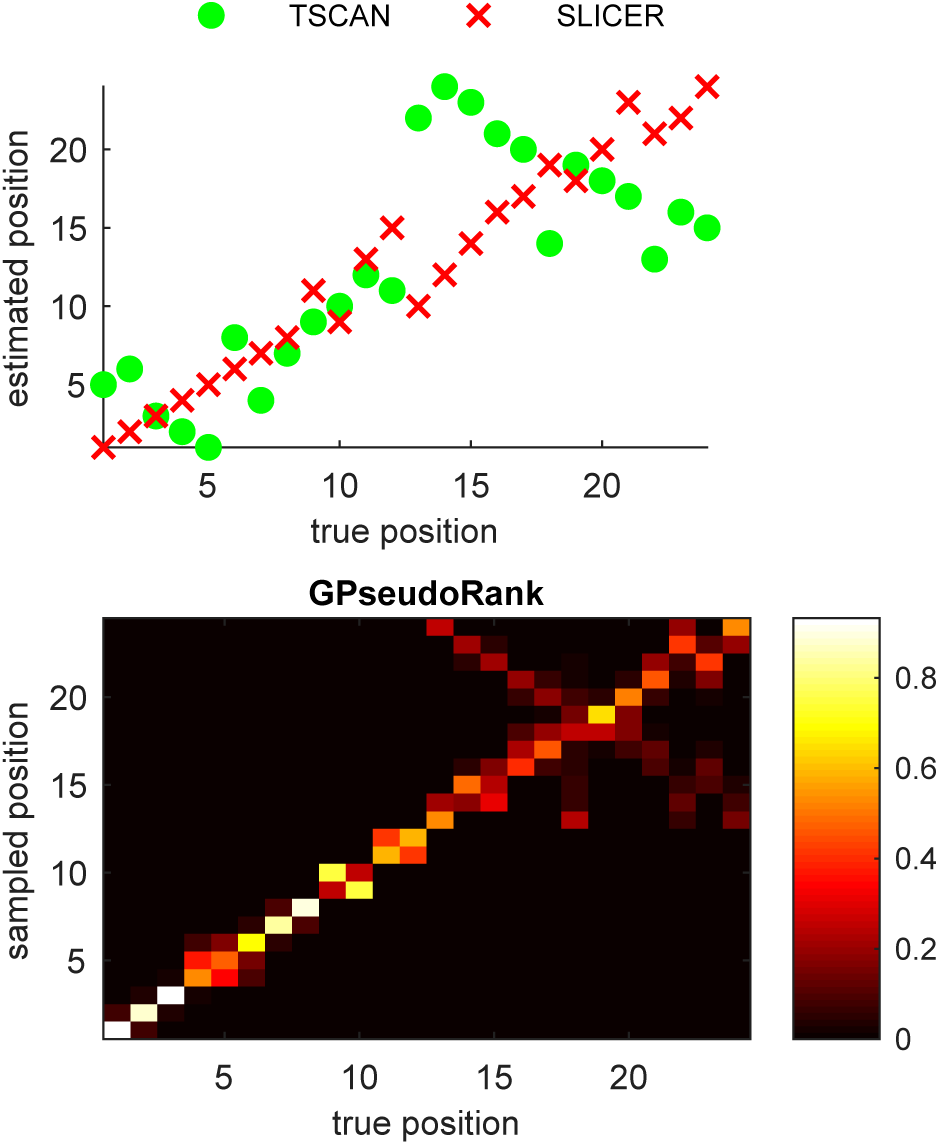
Comparing bi-modal posterior by GPseudoRank to point estimates by TSCAN and SLICER: Windram data. For GPSeudoRank, the matrix illustrates the posterior probabilities of the positions of the cells: the true cell position is along the x-axis, the posterior density is plotted along the y-axis. For TSCAN and SLICER, we plotted along the y-axis the estimated position.

### 3.3 Pseudotemporal uncertainty varies during response to infection

For the scRNA-seq data from Shalek et al. [27], collected at five different capture times, the true cell ordering is unknown. To check convergence of orders the 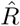-statistic is computed both on the log-likelihood and on the *L*^1^-distances of the permutation of cell positions to an arbitrary reference permutation (Figure 1).

Figure 1 shows that a threshold for the 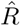-statistic of 1.1 has been reached after 10,000 thinned samples. We therefore discard a burn-in of 5,000 thinned samples at the beginning of each chain, as recommended by Gelman and Shirley [9]. Indeed, by the 1.1 threshold for the 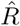 statistic 10,000 thinned samples would have been sufficient for convergence.

Figure 2 demonstrates again the value of providing a posterior distribution for orders, rather than a single estimate: TSCAN and SLICER give different results. The TSCAN result is compatible with the sampled distribution, however, the SLICER result seems to be an outlier. Knowledge of the uncertainty can prevent over-confidence in the results.

Figure 4 illustrates the uncertainty of the pseudotime over the mean pseudotime. To ensure that the inverted U-shape in the amount of uncertainties of the first two capture times at 0h and 1h is not a sampling artifact, cells from these capture times were mixed together for initialising the sampler (that is, capture time information was discarded). On the other hand, despite being separated during initialisation of the sampler, cells from capture times 4h and 6h are completely merged, again indicating that the sample has reached convergence.

**Figure 4:**
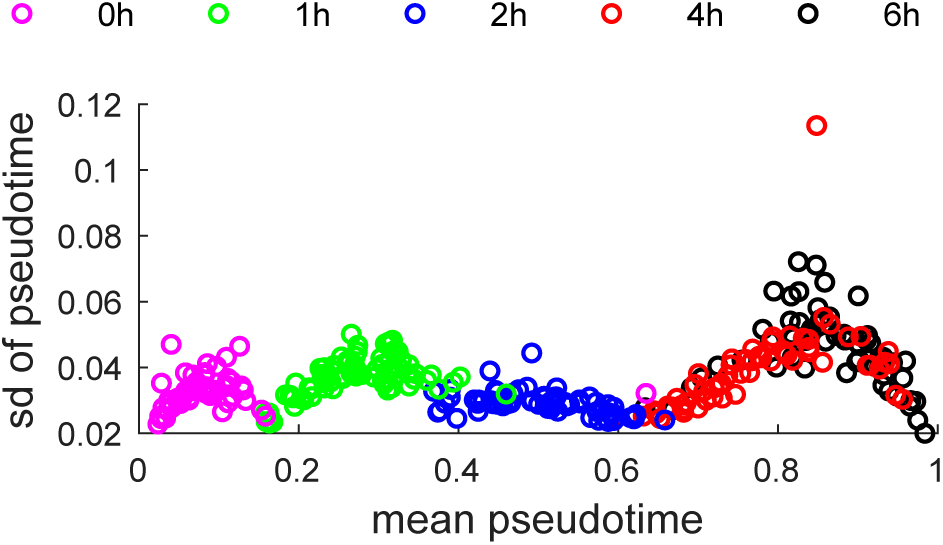
Uncertainty of pseudotime as a function of mean pseudotime. For each cell, the mean pseudotime is plotted along the x-axis, and the respective standard deviation along the y-axis. Cells are coloured by capture time.

Overall uncertainty in the ordering of cells is markedly lower around capture time 2h, when the reaction to the infection has set in, but is not yet complete.

The slight U-shape in the amount of uncertainty for capture times 0h, 1h, and 4h/6h seems to be an experimental batch effect of capturing multiple heterogeneous cells at different time points. Within a batch (or merged batches 4h and 6h) cells which are either lagging behind or slightly ahead in their development are assigned a more specific pseudotime with lower uncertainty behind or ahead of the bulk of cells whose pseudotimes are more interchangeable with higher uncertainty.

GPseudoRank identifies two precocious cells, pointed out in the original analysis by [27], ahead in terms of their response to the stimulus, see Figure 5. Shalek et al. identified a set of genes particularly associated with antiviral response. Ahmed et al. and Reid and Wernisch also used this score to demonstrate that their methods identify two cells at capture time 1h precocious in their antiviral response. Figure 6 shows the average expression of a set of genes associated with antiviral response for each cell. As expected, this antiviral score increases over pseudotime, confirming that the pseudotime assignment captures a biological phenomenon. In contrast to Figure 6, both DeLorean [24] and GrandPrix [1] show considerable edge effects in comparable plots [24, Fig. 4], [1, Fig. 2]. Such edge effects are not biologically motivated and presumably algorithmic artifacts which GPseudoRank is able to avoid by restricting pseudotimes to a finite interval and by using a geodesic mapping.

**Figure 5:**
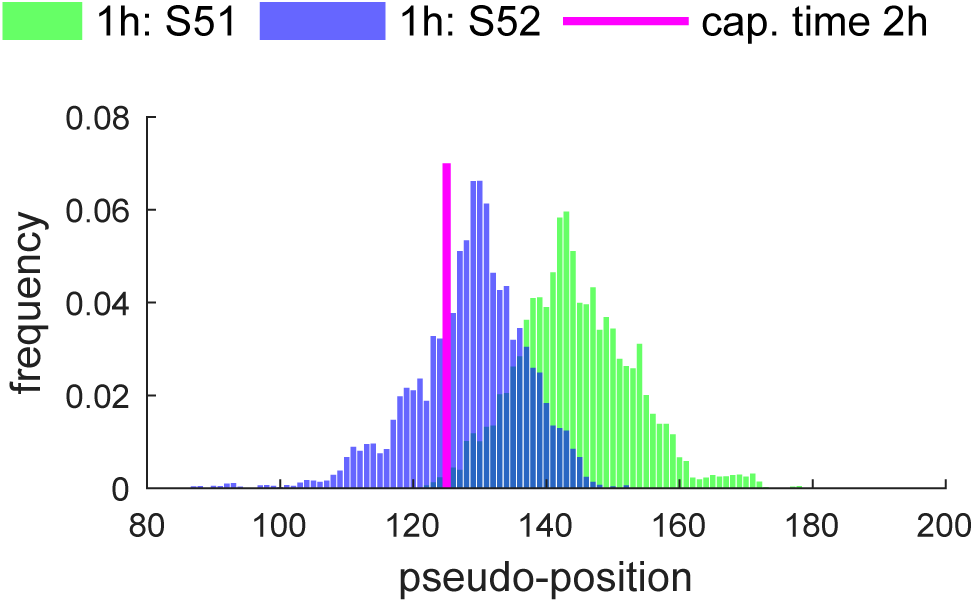
Posterior distribution of cell positions of the precocious cells. For each posterior position of the cells we plot the frequency at which this position occurs among all samples. One random MCMC chain was used. Both of the precocious cells have a high probability of being located within capture time 2h, with S51 likely to be ahead of S52.

**Figure 6:**
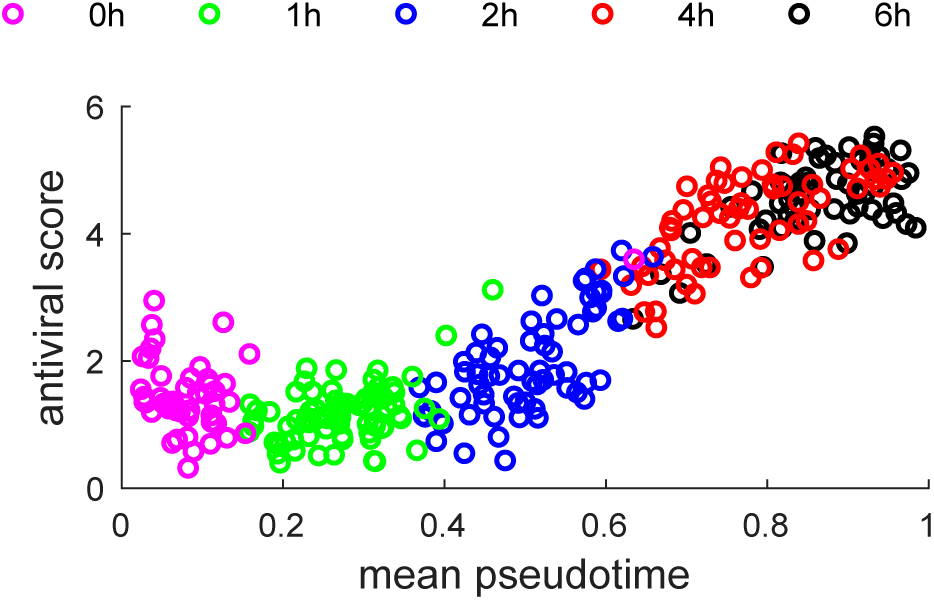
Core antiviral score as a function of mean pseudotime.

## 4 Discussion

GPseudoRank is a new type of Gaussian process latent variable model for pseudotemporal ordering. It samples orderings instead of pseudotimes, with combinatorial proposal moves designed to allow the Metropolis-Hastings sampler to make large changes to permutations and still achieve a high acceptance rate. Figure 2 clearly illustrates the advantage of sampling from a posterior distribution of cell orderings over deriving a single estimate. Although for this data set the true ordering is known, our sampler shows that orders near a second distinct mode are still likely. In fact one such alternative order is returned by a popular algorithm. For data with unknown order, knowledge of the possibility of alternative modes is preferable to the return of just one arbitrary solution.

For the microarray data set we used move 3 only, which reverses whole segments of a permutation, because of its particular suitability for capturing multimodality. We therefore recommend to run two MCMC chains of GPseudoRank in parallel: one with move 3 only, which performs best in case of multi-modality, and a second one with all moves, for faster convergence for less complicated distributions of orders with higher noise levels, as our simulation studies show.

The application to an scRNA-seq data set illustrates another advantage of sampling from the posterior of orderings: the amount of uncertainty about the position of a cell can vary with time. In this case, the uncertainty is lowest in the middle of the process, where the heterogeneity of cells with regard to their progress through the response to the infection is highest. This identifies parts of the process with increased change and higher biological variability compared to technical noise.

The uncertainty of the orders is relevant to any further analysis that models scRNA-seq data in terms of time-series data. This applies, for instance, to any type of network inference where the order of the input time series is relevant, including GP models [20] and vector-autoregressive ones [19]. Alternatively, identifying the regions of the process where the uncertainty of a cell’s position is low can support the selection of suitable cells for the clustering of genes, for example.

Variational inference, which avoids sampling altogether, is considered a computationally efficient if only approximate Bayesian inference alternative to MCMC sampling. However, it turns out that MCMC sampling from discrete permutations in GPseudoRank is efficient enough that its run time is comparable to that of a variational approach: 100,000 iterations for the Windram data take 7min 20s on a single Intel Xeon X5 2.0GHz CPU, compared to about 3 minutes for each initialisation of the variational sampler in DeLorean on one core of an AMD 6174 2.2 Ghz CPU. Similarly, for the scRNA-seq data set, sampling the 100,000 samples shown to be sufficient for convergence takes about 40 minutes, compared to 20 minutes for each initialisation of the variational sampler in DeLorean.

Overall, GPseudoRank offers new insights into biological phenomena and experimental artifacts. It quantifies the amount and variability of uncertainty in single-cell ordering (Figure 4). Assessing the degree of uncertainty enables spotting experimental batch effects created by sampling from a continuous spectrum of developmental stages at only a few capture times. Our approach is also able to identify precocious cells (Figure 5). By combining a geodesic pseudotime mapping with sampling permutations, GPseudoRank also avoids edge effects present in other GP methods for pseudotime ordering (Figure 6).

